# Global assessment of effective population sizes: consistent taxonomic differences in meeting the 50/500 rule

**DOI:** 10.1101/2023.09.22.558974

**Authors:** Shannon H. Clarke, Elizabeth R. Lawrence, Jean-Michel Matte, Brian K. Gallagher, Sarah J. Salisbury, Sozos N. Michaelides, Ramela Koumrouyan, Daniel E. Ruzzante, James W.A. Grant, Dylan J. Fraser

## Abstract

Effective population size (*N_e_*) is a particularly useful metric for conservation as it affects genetic drift, inbreeding and adaptive potential within populations. Current guidelines recommend a minimum *N_e_* of 50 and 500 to avoid short-term inbreeding and to preserve long-term adaptive potential, respectively. However, the extent to which wild populations reach these thresholds globally has not been investigated, nor has the relationship between *N_e_* and human activities. Through a quantitative review, we generated a dataset with 4145 georeferenced *N_e_* estimates from 3576 unique populations, extracted from 712 articles. These data show that certain taxonomic groups are less likely to meet 50/500 thresholds and are disproportionately impacted by human activities; plant, mammal, and amphibian populations had a ≤52% probability of reaching *N̂*_*e*_ = 50 and a <5% probability of reaching *N̂*_*e*_ = 500. Populations listed as being of conservation concern according to the IUCN Red List had a lower *N̂*_*e*_ than unlisted populations, and this relationship held true across all taxonomic groups. *N̂*_*e*_ was reduced in areas with a greater Global Human Footprint, especially for amphibians and mammals, however relationships varied between taxa. We also highlight several considerations for future works estimating *N̂*_*e*_, including the role that gene flow and subpopulation structure plays in the estimation of *N̂*_*e*_ in wild populations, and the need for finer-scale taxonomic analyses. Our findings provide guidance for more specific thresholds based on *N_e_* and help prioritize assessment of populations from taxa most at risk of failing to meet conservation thresholds.

## Introduction

Wild populations face increasing threats from human activities (Bellard et al., 2012; Rosser & Mainka, 2002; Wilson et al., 2016) amidst the Earth’s sixth mass extinction event (Ceballos et al., 2015; Pimm et al., 2014), resulting in a doubling of the number of endangered species over the past decade (IUCN Red List, 2020). Measuring extinction risk is critical to maintaining wild populations and reducing biodiversity loss, by facilitating the appropriate direction of conservation resources. Although demographic and ecological measures are commonly used to evaluate extinction risk, e.g., declines in population size or contractions in range size (Mace et al., 2008), genetic measurements of diversity or inbreeding depression also provide critical, complementary information and are increasingly adopted in conservation assessments (Dunham et al., 1999; Frankham, 2005; Frankham et al., 2014; Hoban, 2020).

A key genetic variable for evaluating extinction risk is the effective population size (*N_e_*) (Frankham et al., 2014; Hoban, 2020; Laikre et al., 2009). Not only does *N_e_* influence the amount of genetic diversity lost through random genetic drift within a population, but it also reflects the level of inbreeding, which can make individuals less fit and thereby generate population declines, making populations with small *N_e_* especially more susceptible to extinction (Bijlsma et al., 2000; Crow & Kimura, 1970; Waples, 2002a). Indeed, previous work has shown that *N_e_* estimates are lower in populations of conservation concern (Palstra & Ruzzante, 2008). The effective size is also closely linked to the demographic history and structure of a population, and furthermore, can be used to project future demographic changes which is of growing interest for conservation practices (Novo et al., 2022; Wang et al., 2016).

*N_e_* is often compared alongside *N_c_* (census size) to evaluate both demographic and genetic changes in managed populations (Palstra & Fraser, 2012). The *N̂*_*e*_ /*N̂*_*c*_ ratio is used in risk assessments or calculation of minimum viable population sizes, and previous reviews have reported means or medians ranging from 0.1 to 0.2 (Frankham, 1995; Palstra & Fraser, 2012; Palstra & Ruzzante, 2008). Thresholds of minimum *N̂*_*e*_ have also been established to ensure wild population persistence (Franklin, 1980). The 50/500 rule recommends a minimum *N̂*_*e*_ of 50 to avoid inbreeding in the short-term, and a minimum *N̂*_*e*_ of 500 to allow sufficient genetic diversity for adaptation in the long-term (Franklin, 1980; Jamieson & Allendorf, 2012). Although the 50/500 criteria is increasingly incorporated into minimum viable population size assessments (e.g., IUCN Red List, Convention on Biological Diversity; Hoban, 2020; Laikre et al., 2020; Mace et al., 2008), the broad applicability of this criteria in conservation is heavily debated (Frankham et al., 2013; Franklin et al., 2014; Jamieson & Allendorf, 2012; Laikre et al., 2021), and some authors recommend increasing the minimum thresholds to 100 and 1000 to maintain populations’ fitness and adaptive potential (Frankham, 2014; García-Dorado, 2015; Jamieson & Allendorf, 2012; Pérez-Pereira et al., 2022).

A global assessment of how well wild populations fulfill 50/500 criteria is timely due to its benefits for forecasting adaptive responses of at-risk populations to environmental change. Furthermore, assessing the spatial positioning of *N_e_* estimates across taxa in relation to human activities could provide significant benefits for conservation planning by contributing to emerging literature on the processes affecting population genetic diversity at broad, macrogenetic scales (e.g., Lawrence & Fraser, 2020; Leigh et al., 2019; Schmidt et al., 2023). Estimates of *N_e_* in wild populations have accumulated rapidly in more than a decade since previous reviews of *N̂*_*e*_ were conducted. These previous reviews lacked adequate sample sizes for taxon-specific analyses and focused heavily on *N_e_* estimates generated using either demographic methods or temporal genetic methods (Frankham, 1995; Hoban, 2020; Palstra & Fraser, 2012; Palstra & Ruzzante, 2008). More recent proliferation of *N_e_* estimates has arisen from the development of single sample approaches to *N_e_* estimation and genome-wide SNPs (single-nucleotide polymorphisms; Barbato et al., 2015; Santiago et al., 2020).

Here, we collate and synthesize global data on single-sample contemporary *N_e_* estimates across taxa using systematic review approaches (e.g., O’Dea et al, 2021), and link these to spatial data obtained through geo-referencing of populations. This approach allows us to: (1) evaluate taxonomic differences in *N̂*_*e*_, *N̂*_*b*_, the likelihood to meet 50/500 thresholds and reassess *N̂*_*e*_ /*N̂*_*c*_ ratios across taxa; (2) compare *N̂*_*e*_ values between threatened and nonthreatened populations (according to the IUCN Red List), and (3) create a global map of *N̂*_*e*_ to assess relationships between *N̂*_*e*_ and spatial patterns in human activities.

We expected taxonomic differences in mean *N̂*_*e*_ and *N̂*_*e*_ /*N̂*_*c*_ ratios because *N_e_* is closely linked with life history characteristics that can vary among taxa. We specifically expected larger *N̂*_*e*_ in groups such as marine fishes, which inhabit large areas with high connectivity between populations (Marandel et al., 2019; Palstra & Ruzzante, 2008); such groups would be most likely to meet 50/500 thresholds as well. We also expected that groups with type III survivorship curves (i.e., high fecundity but low juvenile survival, e.g., marine fishes) would have the lowest *N̂*_*e*_ /*N̂*_*c*_ ratios due to a high variance in reproductive success and relatively large population sizes (Hedrick, 2005; Waples, 2002b).

Similar to what was found by Palstra and Ruzzante (2008), we further expected to find a smaller *N_e_* in populations listed on the IUCN Red List. We also predicted that *N̂*_*e*_ would depend on an interactive effect between IUCN status and taxonomic group, given that some groups, such as amphibians, have a much higher proportion of listed species, and that listed species are expected to have a lower *N̂*_*e*_ (Hoffmann et al., 2010). However, if IUCN status did not interact with taxonomic group to affect *N̂*_*e*_, it would indicate that IUCN status does not explain differences in *N̂*_*e*_ between taxonomic groups.

Finally, we predicted that *N_e_* is negatively correlated with human impact (as measured by the Global Human Footprint; WCS & CIESIN, 2005) because habitat fragmentation, habitat destruction, and pollution can lead to reduced population sizes, reduced connectivity, or may impact breeding, therefore reducing *N̂*_*e*_. The Global Human Footprint is a quantitative assessment of human impact that includes factors such as population density, agriculture, transportation networks, and light pollution (Venter et al., 2016).

## Methods

### Literature search, screening, and data extraction

A primary literature search was conducted using ISI Web of Science Core Collection and any articles that referenced two popular single-sample *N_e_* estimation software packages: LDNe (Waples & Do, 2008), and NeEstimator v2 (Do et al., 2014). The initial search included 4513 articles published up to the search date of May 26, 2020. Articles were screened for relevance in two steps, first based on title and abstract, and then based on the full text. For each step, a consistency check was performed using 100 articles to ensure they were screened consistently between reviewers (n = 6).We required a kappa score (Collaboration for Environmental Evidence, 2020) of ≥ 0.6 in order to proceed with screening of the remaining articles. Articles were screened based on three criteria: (1) Is an estimate of *N_e_* or *N_b_* reported; (2) for a wild animal or plant population; (3) using a single-sample genetic estimation method. Further details on the literature search and article screening are found in the Supplementary Material (Fig. S1). We extracted data from all studies retained after both screening steps (title and abstract; full text). Each line of data entered in the database represents a single estimate from a population. Some populations had multiple estimates over several years, or from different estimation methods (see Table S1), and each of these was entered on a unique row in the database. Data on *N̂*_*e*_, *N̂*_*b*_, or *N̂*_*c*_ were extracted from tables and figures using WebPlotDigitizer software version 4.3 (Rohatgi, 2020). A full list of data extracted is found in Table S2.

### Data Filtering

After the initial data collation, correction, and organization, there was a total of 8971 *N_e_* estimates (Fig. S1). We used regression analyses to compare Ne estimates on the same populations, using different estimation methods (LD, Sibship, and Bayesian), and found that the R^2^ values were very low (R^2^ values of <0.1; Fig. S2 and Fig. S3). Given this inconsistency, and the fact that LD is the most frequently used method in the literature (74% of our database), we proceeded with only using the LD estimates for our analyses. We further filtered the data to remove infinite and negative estimates, and estimates where no sample size was reported or no bias correction (Waples, 2006) was applied (see Fig. S4 for more details). Point estimates with an upper confidence interval of infinity (n = 1358) were larger on average (*N̂*_*e*_ = 478.3, compared to 163.7 and 95.3, for estimates with no CIs or with an upper boundary, respectively).

Nevertheless, we chose to retain point estimates with an upper confidence interval of infinity because accounting for them in the analyses did not alter the main conclusions of our study and would have significantly decreased our sample size (Fig. S5). We also retained estimates from populations that were reintroduced or translocated from a wild source (n = 309), whereas those from captive sources were excluded during article screening (see above). In exploratory analyses, the removal of these data did not influence our results, and many of these populations are relevant to real-world conservation efforts, as reintroductions and translocations are used to re-establish or support small, at-risk populations. We removed estimates based on duplication of markers (keeping estimates generated from SNPs when studies used both SNPs and microsatellites), and duplication of software (keeping estimates from NeEstimator v2 when studies used it alongside LDN_e_). Spatial and temporal replication were addressed with two separate datasets (see Table S3 for more information): the full dataset included spatially and temporally replicated samples, while these two types of replication were removed from the non-replicated dataset. Finally, for all populations included in our final datasets, we manually extracted their protection status according to the IUCN Red List of Threatened Species. Taxa were categorized as “Threatened” (Vulnerable, Endangered, Critically Endangered), “Nonthreatened” (Least Concern, Near Threatened), or “N/A” (Data Deficient, Not Evaluated).

### Mapping and Human Footprint Index (HFI)

All populations were mapped in ArcGIS using the coordinates extracted from articles. The maps were created using a World Behrmann equal area projection. For the summary maps, estimates were grouped into grid cells with an area of 250,000 km^2^ (roughly 500 km x 500 km, but the dimensions of each cell vary due to distortions from the projection). Within each cell, we generated the count and median of *N̂*_*e*_ (using medians here to compare with previous reviews). We used the Global Human Footprint dataset (WCS & CIESIN, 2005) to generate a value of human influence (HFI) for each population at its geographic coordinates. The footprint ranges from zero (no human influence) to 100 (maximum human influence). Values were available in 1 km x 1 km grid cell size and were projected over the point estimates to assign a value of human footprint to each population. The human footprint values were extracted from the map into a spreadsheet to be used for statistical analyses. Not all geographic coordinates had a human footprint value associated with them (i.e., in the oceans and other large bodies of water), so 3299 *N_e_* estimates in our final dataset had an associated footprint value.

### Statistical Analyses

All statistical analyses were performed in R statistical software (version 4.0.5; R Core Team, 2021). To address our main objectives, we constructed generalized linear mixed models (glmms) using the package *glmmTMB* (Magnusson et al., 2017), using backwards model selection with AICc. All models used a gamma distribution with a log link to test whether *N̂*_*e*_, *N̂*_*b*_, and *N̂*_*e*_/*N̂*_*c*_ varied across taxa, with the exception of our 50/500 analysis, which used a binomial distribution relating to whether each threshold of 50 and 500 were met or not (0 if the threshold was not met, and 1 if it was met). All models were run using the replicated dataset, with the exception of the 50/500 analysis, to avoid pseudoreplication in the number of populations that fell below the thresholds. For the 50/500 analysis, we also only used data on *N̂*_*e*_, as the thresholds are specific to *N̂*_*e*_ and not *N̂*_*b*_. For all models, we included taxonomic group as a fixed effect (invertebrates, amphibians, reptiles, freshwater fishes, diadromous fishes, marine fishes, birds, mammals, and plants), as well as marker type (microsatellite, SNP, or other – allozyme) and number of loci (i.e., number of microsatellites or number of SNPs, standardized using the z-score transformation) to control for the effects of these factors. Fishes were separated into three separate groups (diadromous, freshwater, and marine) as these are expected to differ by many ecological characteristics that influence their life history and genetics (DeWoody & Avise, 2000; Martinez et al., 2018). For the IUCN Red List analysis, we included IUCN status and an interaction term between IUCN status and taxonomic group as predictor variables, and we removed any populations that were “N/A” (Data Deficient or Not Evaluated). For the Global Human Footprint analysis, we also used footprint and the interaction between footprint and taxonomic group as predictor variables. For the *N̂*_*e*_/*N̂*_*c*_ analysis, we combined *N̂*_*e*_ and *N̂*_*b*_ data due to a lower sample size of estimates that also included an *N̂*_*c*_ value (253 *N̂*_*b*_ and 286 *N̂*_*e*_ estimates). We also included a dependent variable to indicate whether the ratio was from an *N̂*_*e*_ or an *N̂*_*b*_ estimate. We weighted all models by sample size, using the *weights* function in *glmmTMB*, as larger sample sizes better approximate the true *N̂*_*e*_ of populations (Waples & Do, 2008). Random effects included in each model were the population nested within the study (to account for replication within populations, and any study-level effects), with the exception of the 50/500 analysis, where only study ID was used as a random variable as we were using the nonreplicated dataset. We generated the estimated marginal means for each model by taxonomic group using the *emmeans* package (Lenth, 2018), and plotted the means using *ggplot2* version 3.3.3 (Wickham et al., 2016). These estimated marginal means (emms) account for any differences in the other fixed effects included in the final model. For example, when comparing the emms of *N̂*_*e*_ between taxonomic groups, this would account for any differences in marker type (Table S4). For the Global Human Footprint analysis, we generated values of *N̂*_*e*_ for each taxonomic group at each value of human footprint between 0 and 100, using the function *emmip* within the *emmeans* package, and plotted them using *ggplot2*. We also generated estimates of the intercept and slope for the interaction of *N̂*_*e*_ and HFI for each taxonomic group, using the *emmip* and *emtrends* functions within the *emmeans* package.

## Results

Following systematic review of primary literature and filtering, the full dataset for analysis included 4142 contemporary *N_e_* estimates, 827 effective number of breeder (*N_b_*) estimates, and 534 *N_c_* estimates from 3558 unique populations, extracted from 712 articles. *N̂*_*b*_was included in the review as an estimate of *N_e_* based on a single breeding season. This full dataset included spatially and temporally replicated samples; a second, non-replicated dataset that excluded temporal and spatial replication was also analysed to avoid pseudoreplication (see Table S3 for more information). The non-replicated dataset included 3315 *N_e_* estimates and 343 *N_b_* estimates and was only used for analyses concerning the 50/500 ratio.

Our datasets included studies published from 2006 to 2020, with the number of studies published per year increasing steadily (Fig. S6). Although the year in which populations were sampled ranged from the mid 1900s to 2019, 92% of *N_e_* or *N_b_* estimates came from populations sampled in or after 2000 (Fig. S7). Across all taxa, the median *N̂*_*e*_ was 83.4, and the median *N̂*_*b*_ was 93.0. Freshwater fishes and reptiles had the most and fewest estimates (n = 1390, n = 211), respectively (see Table S5 for more information on sample sizes). Notably, salmonid fishes comprised 31% of our estimates, while only 6% of estimates across all taxa were from populations that had been reintroduced or translocated by humans. Overall, populations sampled were concentrated in Europe and coastal North America, with low representation in Africa and mid-Western Asia (Fig. S8). Median *N̂*_*e*_ values were relatively high in Alaska (100 to >1000), while low values (0-25) were spread out across the globe (Fig. 1).

**Fig. 1:**
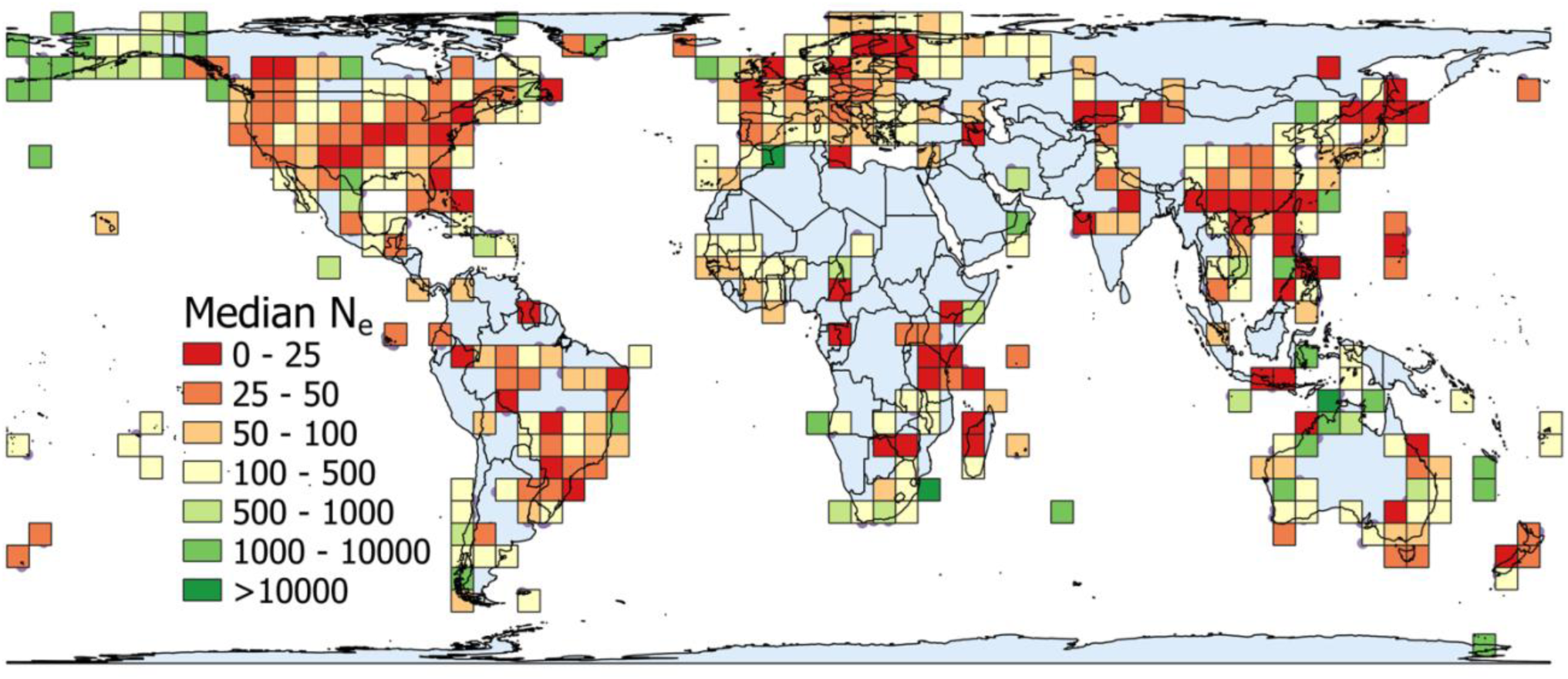
Global map showing the median *N̂*_*e*_ value across all taxonomic groups in each 250,000km^2^ grid cell. Data is projected with the world Behrmann projection.

### Taxonomic Differences in *N̂*_*e*_, *N̂*_*b*_, *N̂*_*e*_/*N̂*_*c*_ and Agreement with the 50/500 Rule

We expected to find taxonomic differences in *N̂*_*e*_ and *N̂*_*b*_, due to their close link to life history characteristics that can vary among taxa. As expected, *N̂*_*e*_ and *N̂*_*b*_ varied by taxonomic group, as revealed from linear mixed models (Table S4). Based on the raw data before modeling, marine fishes had the largest median *N̂*_*e*_ and *N̂*_*b*_ (634.6 and 798.0, respectively; Table 1), while plants had the smallest median *N̂*_*e*_ (35.1), and amphibians had the smallest median *N̂*_*b*_ (52.1). After accounting for spatial and temporal replication and differences in marker type through linear mixed models, marine fishes had a significantly higher *N̂*_*e*_ (estimated marginal mean of 880.2; df = 14, p <0.001) than all other groups except diadromous fishes (*N̂*_*e*_ = 557.0; Table 1). Groups with the lowest *N̂*_*e*_ were plants, amphibians, and mammals (*N̂*_*e*_ = 35.9, 58.3, and 63.8, respectively), while reptiles, invertebrates, freshwater fishes, and birds had intermediate values (Fig. 2-i; Table 1). Comparing *N̂*_*b*_ values across taxonomic groups, marine fishes again had the largest mean (*N̂*_*b*_ = 1175.3), however they were statistically indifferent from several other groups (plants, invertebrates, birds, and diadromous fishes; Fig. S9). Groups with the lowest *N̂*_*b*_ were amphibians, reptiles, and mammals (*N̂*_*b*_ = 48.2, 50.9, and 60.1, respectively). Marker type used to estimate *N_e_* was also an explanatory variable in the most parsimonious models (both p < 0.001; Table S4); SNPs had a higher median *N̂*_*e*_ compared to microsatellites (*N̂*_*e*_ = 174.5 vs. 75.9), but microsatellites generated higher estimates than SNPs on average in the model (*N̂*_*e*_ = 129 vs. 104, respectively, and *N̂*_*b*_ = 166 vs. 109, respectively). This is likely due to the smaller number of SNP estimates overall, and the non-random targeting of species for studies using SNPs (see discussion).

**Fig. 2:**
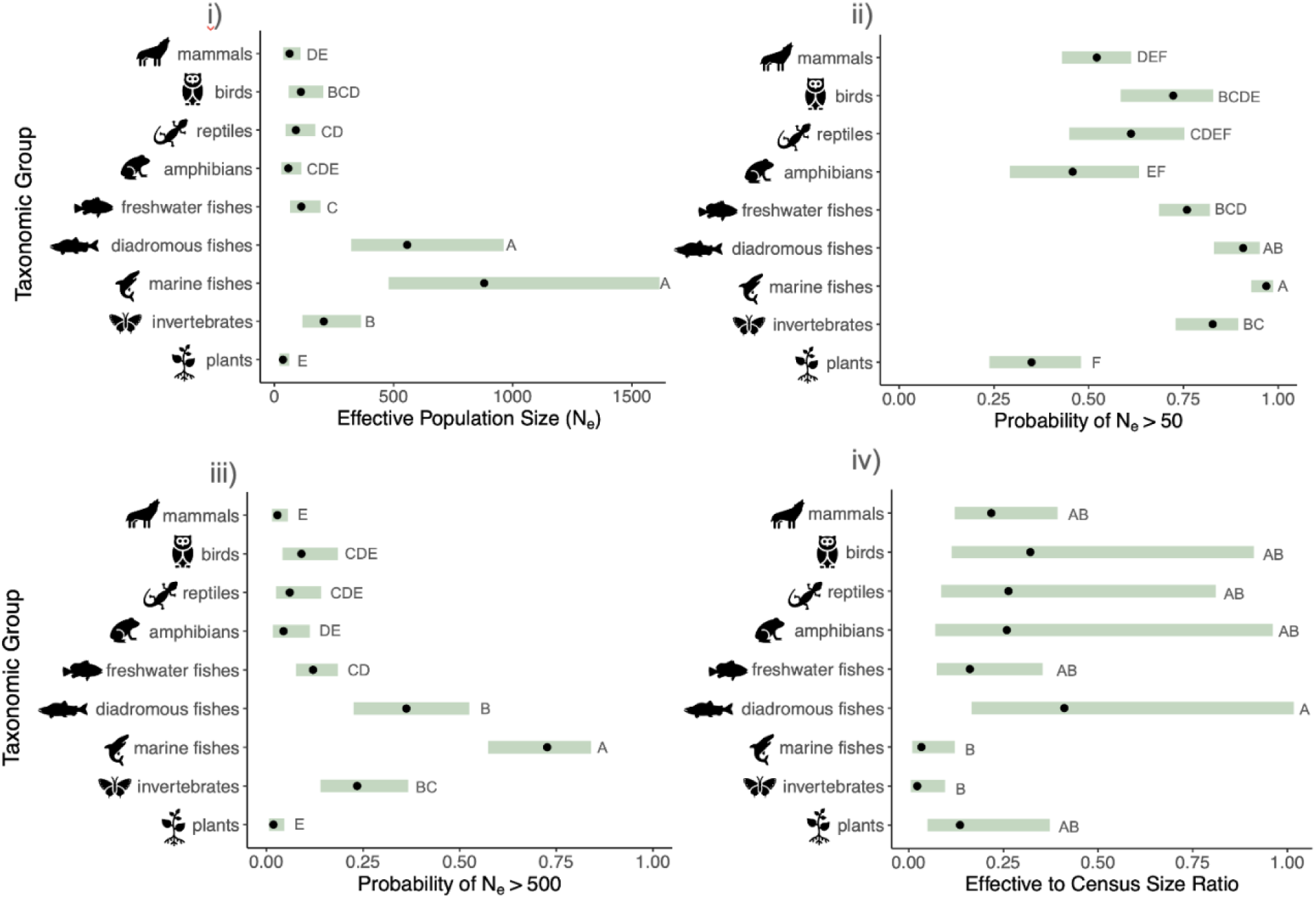
i) Mean effective population size (*N̂*_*e*_) across taxonomic groups, accounting for differences in marker type. ii) Probability of meeting the lower threshold of 50/500 rule (*N̂*_*e*_ = 50). iii) Probability of meeting the upper threshold of 50/500 rule (*N̂*_*e*_ = 500). Shaded green bars represent the 95% confidence intervals. Groups with no shared letters are statistically different (*α* = 0.05) from one another. iv) Effective to census size ratio (*N̂*_*e*_ /*N̂*_*c*_ and *N̂*_*b*_/*N̂*_*c*_) across taxonomic groups, accounting for differences between *N̂*_*e*_ and *N̂*_*b*_.

**Table 1.** Summary statistics between and across taxonomic groups for N̂_e_ and HFI analyses, including sample sizes, medians, model estimated means, and slopes.

| Taxonomic Group | $\hat{N}_e$ Analysis | | | HFI Analysis | |
| --- | --- | --- | --- | --- | --- |
| | Sample Size | Median $\hat{N}_e$ | Model-Estimated Means (± SE) | Sample Size | Slope of HFI * Taxonomic Group |
| Amphibians | 369 | 43.3 | $58.3 \pm 20.0$ | 339 | -0.0238 |
| Diadromous Fishes | 686 | 211.5 | $557.0 \pm 155.0$ | 506 | 0.0059 |
| Birds | 204 | 77.8 | $111.2 \pm 34.8$ | 165 | -0.0200 |
| Freshwater Fishes | 1113 | 80.8 | $113.0 \pm 31.1$ | 1045 | -0.0127 |
| Invertebrates | 321 | 135.6 | $207.1 \pm 59.4$ | 216 | -0.0178 |
| Mammals | 552 | 47.2 | $63.8 \pm 17.8$ | 477 | -0.0185 |
| Marine Fishes | 248 | 634.6 | $880.2 \pm 272.7$ | 22 | -0.0047 |
| Plants | 439 | 35.1 | $35.9 \pm 10.4$ | 387 | 0.0199 |
| Reptiles | 201 | 72.3 | $90.1 \pm 29.5$ | 132 | -0.0092 |
| **Overall** | **4145** | **83.9** | **N/A** | **3299** | **-0.024** |

In relation to the 50/500 rule, 38.7% of all populations (1282 of 3315) fell below the threshold of 50, and 85.2% of populations (2825 of 3315) fell below 500 (Fig. 2-ii, Fig. 2-iii). Marine fishes were most likely to meet both the 50 and 500 thresholds (probabilities of 0.97 and 0.73, respectively), while all other groups were likely to meet only the 50 and not the 500 threshold or were not likely to meet either threshold (Fig. 2-ii, 2iii). Plants, amphibians, and mammals were least likely to meet either threshold, with probabilities of 0.35, 0.46, and 0.52, respectively, of meeting the lower threshold. For the analysis assessing the number of populations meeting the threshold of 500, marker type used to estimate *N_e_* was also an explanatory variable in the most parsimonious models (p < 0.003, Table S4); *N_e_* estimates generated using SNPs were more likely to reach the threshold than those estimates generated using microsatellites (probability = 0.17 and 0.07, respectively), likely due to the higher median estimate for SNPs.

The ratio between *N̂*_*e*_ or *N̂*_*b*_ and *N̂*_*c*_ also varied between taxa (Table S4), with *N̂*_*b*_/*N̂*_*c*_ ratios smaller on average than *N̂*_*e*_ /*N̂*_*c*_ (0.123 and 0.174, respectively). Diadromous fishes had the highest ratio on average (0.41), while invertebrates and marine fishes had the lowest ratios (0.02 and 0.03, respectively, Fig. 2-iv). The median ratio from the raw data across all taxa was 0.24, with invertebrates having the lowest median (0.09), and birds having the highest median (0.48; Table S6).

### IUCN Red List Status and *N̂*_*e*_

In line with previous work, we expected to find that populations listed on the IUCN red list had a smaller *N̂*_*e*_, and that there would be an interaction between IUCN status and taxonomic group. Overall, 19.1% of populations in our database were listed as threatened, though this ranged from 10.9% in diadromous fishes to 48.8% in reptiles (Table S7). *N̂*_*e*_ differed between threatened and nonthreatened populations (p < 0.02, df = 8, χ^2^ = 1081.7; Table S4), with threatened populations having a smaller *N̂*_*e*_ than nonthreatened (98.7 and 129.4, respectively). However, we found no interactive effect of IUCN status and taxonomic group on *N̂*_*e*_ (Table S4).

### *N̂*_*e*_ and Global Human Footprint (HFI)

Based on the negative influence of human activities such as habitat destruction and pollution on wild populations, we expected to find a negative correlation between *N̂*_*e*_ and human footprint. Across all taxonomic groups, *N̂*_*e*_ declined with increasing HFI (p < 0.001, slope = - 0.024, df = 23, χ^2^ = 1024.4; Table S4), with seven of nine groups showing this negative trend (Fig. 3, Table 1). The strength of relationship between *N̂*_*e*_ and human footprint also varied between groups. For example, amphibians experienced a 90.8% decrease in *N̂*_*e*_ (from 135.7 to 12.5) as human footprint increased from 0 to 100, whereas marine fishes experienced a 37.3% decrease (from 772.0 to 483.9). However, two taxonomic groups had a positive relationship with human footprint (Fig. 3): increasing from a human footprint of 0 to 100 was associated with an increase in *N̂*_*e*_ from 235.8 to 426.8 in diadromous fishes (81.0%), and an increase from 19.1 to 139.6 in plants (631.7%).

**Fig. 3:**
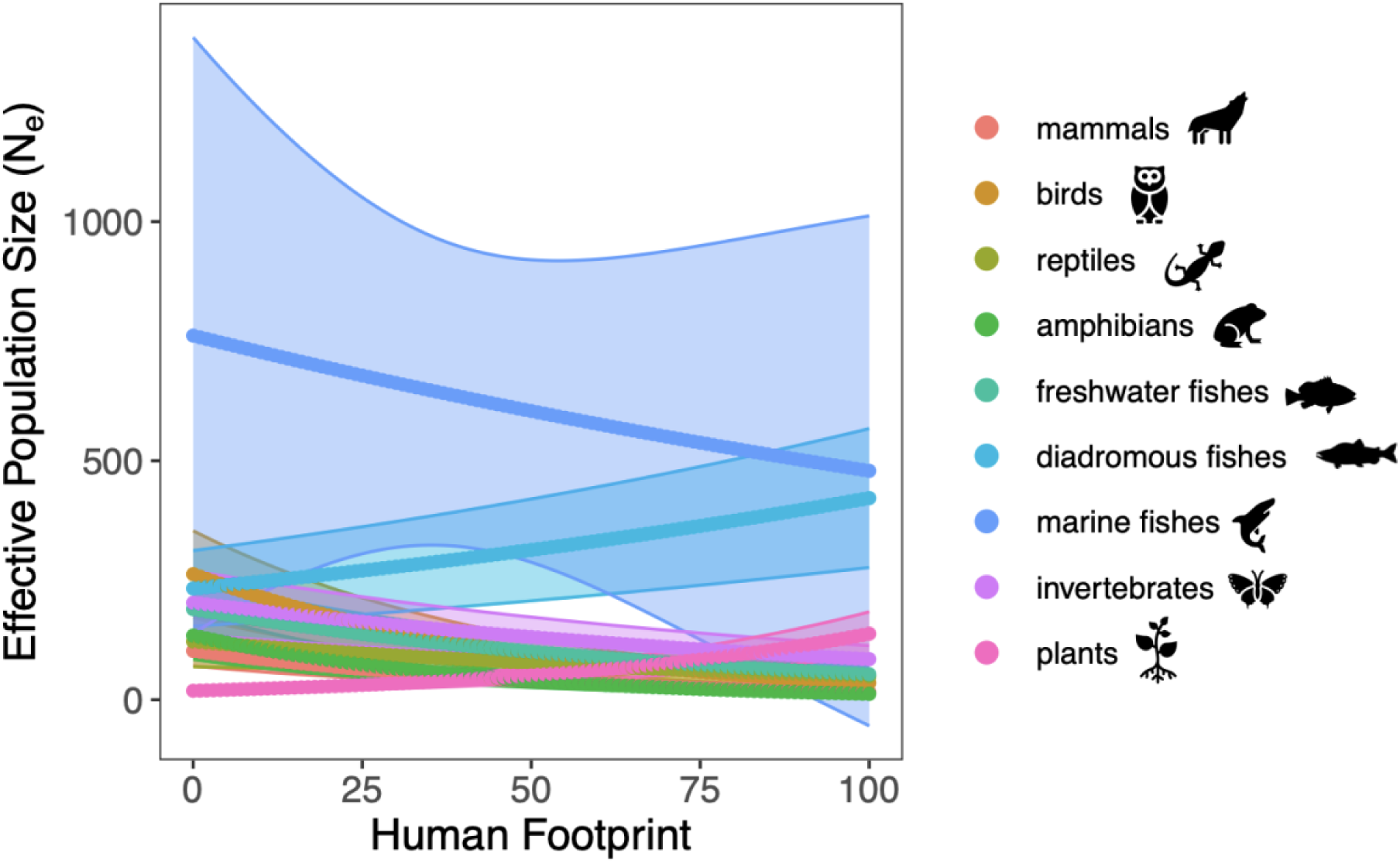
Relationship between effective population size (*N̂*_*e*_) and human footprint across taxonomic groups. Shaded areas represent 95% confidence intervals.

## Discussion

Assessing *N̂*_*e*_ values across taxa on a global scale is essential to prioritise resource allocation for conservation goals and help forecast adaptive responses of populations to future stressors (Hoban et al., 2021). Here, we found that there are consistent differences in *N̂*_*e*_, *N̂*_*b*_, and *N̂*_*e*_ / *N̂*_*c*_ ratios across broad taxonomic groups and that many populations do not meet the 50/500 rule. The probability of meeting 50/500 thresholds also differed among taxa, with amphibians, mammals, and plants being least likely to meet either threshold. Populations listed as threatened, according to the IUCN Red List, had a lower *N̂*_*e*_ than unlisted populations, and this relationship was consistent across all taxonomic groups. While perhaps unsurprisingly, *N̂*_*e*_ decreased in general as Global Human Footprint increased, we also found that this relationship differed between taxonomic groups, with two groups (diadromous fishes and plants) exhibiting a positive correlation with human footprint. These results suggest that previous reviews reporting average or median *N̂*_*e*_ or *N̂*_*e*_ /*N̂*_*c*_ across multiple taxonomic groups may not properly represent patterns within taxa. Our synthesis may help in prioritizing taxa that are unlikely to meet conservation thresholds or that are most impacted by human activities and provides a benchmark for adjusting thresholds based on taxonomic group.

We found large taxonomic differences in *N̂*_*e*_ and *N̂*_*b*_, with marine fishes having the largest *N̂*_*e*_, and amphibians, mammals, and plants having the smallest *N̂*_*e*_. Large *N̂*_*e*_ in populations of marine species was expected, considering they can inhabit large areas, supporting census sizes of thousands to billions of individuals, with high levels of gene flow between connected populations (Hare et al., 2011; Marandel et al., 2019; Palstra & Ruzzante, 2008). Amphibians and mammals, conversely, may be expected to exhibit low *N_e_* values due to their generally small and declining populations, fragmented habitats, and higher risk for extinction compared to other vertebrates. Amphibians are particularly threatened by habitat loss and pollution (Ripple et al., 2017), have the highest proportion of threatened species out of all vertebrates (41% on the IUCN Red List; Hoffmann et al., 2010) and are specifically at risk from the pathogen *Bd*, which has been found in 42% of amphibian species (Olson et al., 2013). Mammals have 25% of assessed species listed on the IUCN Red List (Hoffmann et al., 2010), and terrestrial mammals (which made up >91% of our mammal *N_e_* estimates) are threatened by habitat fragmentation and loss (Crooks et al., 2017). Interestingly, however, while mammals had a relatively high percentage of threatened populations in our dataset (27.2%; Table S7), the amphibian dataset only had 15.3% of populations listed as threatened. Additionally, we found no interaction between IUCN red list and taxonomic group in explaining *N̂*_*e*_. Therefore, lower *N̂*_*e*_ in amphibians and mammals may reflect their generally small and declining populations and higher risks for extinction, but these lower estimates cannot be explained by IUCN listing status alone.

The taxonomic differences in *N̂*_*b*_ generally exhibited the same patterns as *N̂*_*e*_, however confidence intervals were much wider, and statistical differences between groups were fewer, likely due to reduced sample sizes (Table S5). Plants had the largest difference between *N̂*_*e*_ and *N̂*_*b*_ (*N̂*_*e*_ = 36, vs *N̂*_*b*_ = 336), but this may be due to a very low sample size for *N̂*_*b*_ (n = 5). Several explanations could account for the low *N_e_* values in plants, including alternate reproductive strategies which may reduce *N̂*_*e*_ (e.g. selfing or cloning; Orive, 1993; Wright et al., 2013), or declining populations due to climate change, human development, or overharvesting (39% of assessed vascular plant species are threatened with extinction (Nic Lughadha et al., 2020)).

Marine fishes were, as expected, most likely to reach 50/500 thresholds, while amphibians, mammals, and plants were least likely. Most populations across taxa (85.2%) fell below the upper threshold of 500, while 38.7% of populations fell below 50. These results show a much larger proportion of populations are potentially at risk of inbreeding in the short-term (i.e., *N̂*_*e*_ < 50) compared to a 2008 review that reported only ∼8% below 50 (∼70% of populations fell below 500; Palstra & Ruzzante, 2008). In fact, using newer 100/1000 recommendations (Frankham et al., 2014), ∼56% and >90% of populations in this review would fall below *N̂*_*e*_ of 100 and 1000, respectively. One important factor to note, however, is that this review focuses solely on estimates using the Linkage Disequilibrium method, which measures *N̂*_*eLD*_, and can underestimate inbreeding effective size (*N̂*_*eI*_) or additive genetic variance effective size (*N̂*_*eAV*_), which are relevant for the 50/500 rule (Ryman et al., 2019). This underestimation is especially true when sampling from subdivided populations affected by migration, a factor that was not addressed in much of the primary literature upon which this study is based. Due to this, and other potential sources of bias in our dataset (discussed further below), the proportion of populations falling below the 50/500 thresholds is likely overestimated, however the comparisons between groups should hold true. Furthermore, life history traits and other biological differences are expected to play a role in determining whether a population meets these thresholds (e.g., in marine fishes). Moving forward with applying the 50/500 rule to conservation, there is a need to disentangle whether the differences between taxonomic groups are due to declining populations, or if some groups can naturally exist at higher or lower *N̂*_*e*_.

The mean and median *N̂*_*e*_ /*N̂*_*c*_ ratios across taxa were in line with earlier reviews (Hoban, 2020; Palstra & Fraser, 2012; Palstra & Ruzzante, 2008), but varied widely between groups; after accounting for replication and ratio type (i.e. *N̂*_*e*_/*N̂*_*c*_ vs. *N̂*_*b*_/*N̂*_*c*_), marine fishes and invertebrates had smaller ratios on average, (<0.05) and diadromous fishes had the largest ratio. Taxonomic differences in *N̂*_*e*_ /*N̂*_*c*_ were expected because of how life history traits influence the relationship between *N̂*_*e*_ and *N̂*_*c*_ (Waples et al., 2013). The occurrence of low *N̂*_*e*_ /*N̂*_*c*_ ratios in marine fish populations (Hauser et al., 2002; Hoarau et al., 2005; Turner et al., 2002) and other species with Type III survivorship curves (Hedgecock, 1994; Hedrick, 2005; Waples, 2002b) is associated with high fecundities and high juvenile mortalities. Interestingly, invertebrates also had a low *N̂*_*e*_/*N̂*_*c*_ ratio, though 50% of these ratio estimates came from marine molluscs which are broadcast spawners with Type III attributes (Ramirez Llodra, 2002). The high *N̂*_*e*_ /*N̂*_*c*_ ratio of diadromous fishes could be in part due to the high proportion of Atlantic salmon estimates comprising this group (62%). In Atlantic salmon, small non-diadromous males take part in mating and therefore contribute to *N̂*_*e*_, but are often not counted as part of *N̂*_*c*_, resulting in an over-estimation of *N̂*_*e*_/*N̂*_*c*_ (Johnstone et al., 2013; Perrier et al., 2014; Saura et al., 2008). Overall, we caution against generalizing *N̂*_*e*_/*N̂*_*c*_ ratios across taxa with existing data. Data for some groups are still very limited (marine fishes and invertebrates had the smallest sample sizes in this synthesis; n = 9 and n = 6, respectively; Table S5). Furthermore, low *N̂*_*e*_ /*N̂*_*c*_ ratios in marine species could be biased downwards due to an under-estimation of true *N̂*_*e*_ in especially large populations which has been detected applying the linkage disequilibrium (LD) method (Waples, 2016). Lastly, many studies did not report uncertainty around *N_c_* estimates, a limitation which has been previously noted (Palstra & Fraser, 2012). Removing these estimates from this synthesis would have reduced sample sizes from n = 537 to n = 167. Instead, we chose to retain all *N̂*_*c*_ estimates to maximise our sample size, with the caveat that some *N̂*_*c*_ estimates may be less accurate than others. It remains extremely important for authors to report uncertainty around both *N̂*_*e*_ and *N̂*_*c*_, particularly because *N̂*_*e*_/*N̂*_*c*_ ratios can vary substantially among populations *within* some species (e.g., Bernos & Fraser, 2016; Ferchaud et al., 2016).

In line with previous reviews, we found that populations listed as a conservation concern had a lower *N̂*_*e*_ than their unlisted counterparts (98.7 vs. 129.4). Interestingly, however, we did not find any interaction between taxonomic group and IUCN status, indicating that the IUCN Red List designation did not explain the differences in *N̂*_*e*_ between taxonomic groups (i.e., that some groups had lower *N̂*_*e*_ due to having a higher proportion of populations listed on the IUCN). This observation generally holds true when looking at the percentage of populations within each taxonomic group that were listed on the IUCN. Amphibians and plants, which had the lowest mean and median *N̂*_*e*_ only had 15.3% and 22.3% of their populations listed on the IUCN, whereas marine fishes, which had by far the largest mean and median *N̂*_*e*_ had 33.8% of its populations listed (Table S7). It is important to note, however, that the IUCN Red List is a global listing organization that may lack some of the nuance required when assessing population-level differences. Collecting population-specific listing criteria from regional or national organizations was not feasible for this project but could reveal finer-scale trends that could help explain these results further. Regional data may also increase sample sizes for groups such as freshwater fishes or invertebrates, many of which were not assessed, or categorized as “Data Deficient” (41.5% of freshwater fishes and 65.1% of invertebrates).

We found support for the prediction of reduced *N̂*_*e*_ across populations in regions with greater human footprint, however this relationship varied between taxonomic groups in both strength and direction. The human footprint gives a measure of human land use, population density, and infrastructure (e.g., built environments and roads) (WCS & CIESIN, 2005), all of which can contribute to habitat degradation, fragmentation and/or loss, and may result in reduced connectivity, population sizes, and *N_e_*. Several studies have shown decreases in *N̂*_*e*_ associated with anthropogenic habitat fragmentation in single species of different taxa (Alò & Turner, 2005; Browne & Karubian, 2018; Keller et al., 2005; Sumner et al., 2004). The overall relationship with *N̂*_*e*_ and human footprint, while inversely correlated, was relatively weak (with a coefficient of −0.024), likely due to the difficulty in generalizing across a wide variety of species. Indeed, some mammal species have been shown to benefit from human activities and urbanization, leading to more connectivity and a larger habitable area, which could lead to increased *N̂*_*e*_ (Habrich et al., 2021).

Within taxonomic groups, amphibians and mammals had the largest decreases in *N̂*_*e*_ as footprint increased from 0 to 100 (90.8% and 84.3% decreases, respectively). Among mammal populations, human footprint is positively associated with extinction risk, anthropogenic mortality (Di Marco et al., 2018; Hill et al., 2020) and negatively associated with *N̂*_*e*_ (Schmidt et al., 2021) on a continental scale, yet human footprint has been positively associated with amphibian *N̂*_*e*_ (Schmidt & Garroway, 2021) on the same scale. Interestingly, for plants and diadromous fishes, *N̂*_*e*_ was positively correlated with human footprint. In plants, this relationship may be due to positive influences of human activities on dispersal and gene flow, like transportation networks or agricultural development (Auffret et al., 2014; Bullock & Pufal, 2020). For diadromous fishes, it may be that humans preferentially build infrastructure and settlements around larger waterbodies that sustain larger populations of commercially valuable fish (which would also have larger *N̂*_*e*_), such that there is a covariance between larger populations of diadromous fish and increased human footprint.

We have performed a timely review of the *N_e_* literature, with strict literature filtering criteria, georeferencing, and global representation. However, *N_e_* estimates here may be biased due to several factors. First, the Linkage Disequilibrium (LD) method has been reported to underestimate *N̂*_*e*_, especially in populations where the sample size is smaller than the true *N*_*e*_, and in populations with a very large *N*_*e*_ (Waples & Do, 2010). However, with the exception of the 50/500 rule (which objectively compares values to a threshold), the analyses in this study focused on comparisons between groups, all of whom were estimated using the LD method. Furthermore, the LD method is by far the most commonly used in the literature (as evidenced by it comprising almost three quarters of our dataset), and this review aimed to summarize estimates in the literature and can be used as a reference for research using the LD method in the future (even with its limitations). The second potential downward bias was that we removed any infinite or negative estimates, as these could not be incorporated into our analytical results. Some of these removed populations could have been large, and/or the original study sample size used was insufficient to generate a point-estimate of *N_e_* (Do et al., 2014). We suspect, however, that trends observed between taxa would remain similar, as most groups had a similar proportion of infinite values (10-30%). Another caveat of the LD method is that it assumes a closed population and can underestimate *N̂*_*e*_ in populations subject to migration (Ryman et al., 2019). Many of the studies from which we extracted estimates did not address whether populations were open or closed, nor did they attempt to estimate genetic neighbourhood size or metapopulation *N̂*_*e*_. Although we acknowledge the importance of addressing migration in *N*_*e*_ estimation, genetic neighbourhood size or metapopulation *N̂*_*e*_ is not always simple to quantify, and incorporating migration does not always result in a larger *N̂*_*e*_ for subpopulations (Fraser et al., 2007; Gomez-Uchida et al., 2013; Neel et al., 2013; Palstra & Ruzzante, 2011). The problem of addressing migration also holds true for other types and methods of estimating *N̂*_*e*_, such as the temporal method, which estimates variance effective size (Ryman et al., 2023). We encourage future works to consider migration and such additional considerations for the estimation of *N̂*_*e*_, especially when comparing their estimates with the 50/500 rule, to ensure they are not overestimating risk to populations. Lastly, there may be sampling bias associated with which populations are studied or reported in the literature. At-risk populations with small or declining *N̂*_*e*_ and *N̂*_*c*_ may be more commonly studied due to funding from governmental agencies (Mahoney, 2009) or to more interest in published results for threatened species. Overall, although downward biases in reported *N_e_* estimates may exist, we suspect the taxonomic differences still hold true, and these results still provide valuable information on how the 50/500 rule may not be applicable to all taxa.

### Conclusions and recommendations

Our study revealed important differences across broad taxonomic groups in estimates of *N_e_* and in the relationship of these groups with the 50/500 rule and with the Global Human Footprint. Most wild populations did not reach both critical conservation thresholds, and generally *N̂*_*e*_ was negatively associated with human footprint. We identified certain taxonomic groups (i.e., amphibians and mammals) most at risk of falling below these thresholds, potentially due to the negative impacts of human activities. These results can help guide future recommendations for species listing criteria or prioritize assessment of certain taxa most at risk of falling below conservation thresholds. However, these results also raise important questions to be addressed in future works:

(1) Are there finer-scale differences in *N̂*_*e*_ between taxonomic groups (e.g., family, genus, or species)? Studies on the relationship between genetic diversity and urbanization or anthropogenic fragmentation have shown that individual species or genera can vary in the strength and direction of their responses (Habrich et al., 2021; Schmidt et al., 2021). We limited our scope to broad taxonomic groupings of populations, but data exist from certain genera or species (e.g., salmonid fishes) to generate much finer-scale estimates of *N_e_* and further explore the influence of human footprint on *N_e_* within species.
(2) How should the 50/500 or 100/1000 rule be applied to varying taxa? These rules are generalizations based on shared responses across taxa and predictions based on inbreeding depression, fitness, and population size (Frankham et al., 2014). Our synthesis reveals that some taxa may exist at much larger *N̂*_*e*_ than others (i.e., marine fishes), and therefore the 50 and 500 thresholds may be less appropriate for them (with the caveat that *N̂*_*eLD*_ is properly employed under the assumption of a closed population, or genetic neighbourhood size or metapopulation *N̂*_*e*_ is estimated instead; Neel et al., 2013). Future work should focus on disentangling whether the difference seen here between taxonomic groups is due to declining populations (e.g., in amphibians and mammals), or to natural biological/life history trait differences that affect the spatial scale of population differentiation, thereby impacting *N_e_* values. Specifically, focusing on temporal trends and demographic histories can help reveal whether wild populations that exhibit low *N̂*_*e*_ have always done so, or whether the *N̂*_*e*_ has been declining through time.

## Acknowledgements

This work was supported by a Natural Sciences and Engineering Research Council (NSERC) Strategic Project Grant, and Concordia University Research Chair through D. Fraser. S. Clarke was supported by an NSERC Canada Graduate Scholarship (CGS-M), along with a collaborative partnership project award through the Interuniversity Research Group in Limnology (GRIL). D. Ruzzante was supported by an NSERC Discovery grant (RGPIN-2019-04679).

## Author Contributions

**Shannon H. Clarke**: Conceptualization, Methodology, Investigation, Data Curation, Formal Analysis, Visualization, Writing – Original Draft. **Elizabeth R. Lawrence**: Investigation, Formal Analysis, Visualization, Writing – Review & Editing. **Jean-Michel Matte**: Investigation, Formal Analysis, Visualization, Writing – Review & Editing. **Brian K. Gallagher**: Investigation, Formal Analysis, Writing – Review & Editing. **Sarah J. Salisbury**: Investigation, Writing – Review & Editing. **Sozos N. Michaelides**: Investigation, Writing – Review & Editing. **Ramela Koumrouyan**: Investigation, Writing – Review & Editing. **Daniel E. Ruzzante**: Conceptualization, Writing – Review & Editing, Funding Acquisition. **James W. A. Grant**: Supervision, Writing – Review & Editing, Funding Acquisition. **Dylan J. Fraser**: Conceptualization, Supervision, Writing – Review & Editing, Funding Acquisition.

## Data Availability Statement

The data associated with this manuscript will be available on Dryad upon acceptance of the paper for publication

