## Supplemental Material for "Global assessment of effective population sizes: consistent taxonomic differences in meeting the 50/500 rule"

**Supplementary Information**

*Literature Search, Screening, and Data Extraction*

A primary literature search was conducted using ISI Web of Science Core Collection with the following search terms: (“effective population size” OR “effective size” OR *N_e_* OR “effective number of breeders” OR *N_b_*) AND (microsatellite* OR allozyme* OR SSR* OR SNP* OR “single nucleotide polymorphism” OR temporal method*). Effective number of breeders (*N_b_*) was included in the search because it is an estimate of effective size based on a single breeding season. An additional search was done to include articles that referenced two popular single-sample *N_e_* estimation software packages: LDNe, and NeEstimator v2. Article information was exported into MS Excel, and duplicates were removed, resulting in a total of 4513 articles. The database includes articles published up to the search date of May 26, 2020.

Articles were screened for relevance in two steps, first based on title and abstract, and then based on the full text. If article relevance was unclear at the first step, the article was retained and reviewed further at the second step. For each step, a consistency check was performed to ensure the articles were screened consistently between reviewers (n = 6). All reviewers were given a subset of articles to screen (100 for title and abstract and 20 for full text), and their consistency was determined using a Kappa test, where a score of 0 indicates the reviewers agreed no better than expected by chance, and a score of 1 indicates perfect agreement (Collaboration for Environmental Evidence, 2020). A Kappa score of ≥ 0.6 was necessary in order to proceed with screening.

Articles were screened based on three criteria: (1) Is an estimate of *N_e_* or *N_b_* reported; (2) for a wild animal or plant population; (3) using a single-sample genetic estimation method? For criterion (1), only primary literature was included, or where authors obtained DNA sequence information from previous studies and generated a novel estimate of *N_e_* or *N_b_*. For criterion (2), populations that were wild, but supplemented from captive breeding or hatchery releases, were not included, except where hatchery introgression was measured to be very low (<5%; an intermediate between the two options discussed by Allendorf et al. (2004) and Vähä and Primmer (2005)). Populations that were introduced, reintroduced, or translocated were only included if from a wild source. Additionally, estimates of *N_e_* for human populations, parasites, and disease vectors were excluded as they did not fit within the scope of the study. To define populations, reviewers primarily used *F*_ST_ (significance, or values >0.02) and STRUCTURE clusters (Waples & Gaggiotti, 2006). If either method was unavailable, then reviewers relied on other structuring programs (e.g., BAPS, GENELAND), or inferred population structure based on previous literature, or geographic isolation. Studies where the population definition was imprecise or where *N_e_* was estimated above the population level (e.g., if an overall estimate of *N_e_* is provided but *F*_ST_ and STRUCTURE indicate spatial structure) were not included. Studies where *N_e_* was estimated multiple times within a population were included, but were all indexed by the same population ID. For criterion (3), we focused on five main *N_e_* estimation methods, implemented in four software applications (see Table S1). We also restricted our estimation methods to only those that used microsatellite, SNP, or allozyme data.

We extracted data from all studies retained after both screening steps and a final consistency check was conducted, involving all reviewers (n = 6) extracting data from the same 20 articles and comparing the information. Any inconsistencies were discussed, and the data extraction protocol was modified to address them. Each line of data entered in the database represents a single estimate from a population. Some populations had multiple estimates over several years, or from different estimation methods, and each of these was entered on a unique row in the database. Data on $\hat{N}_{e}$, $\hat{N}_{b}$, or $\hat{N}_{c}$ were extracted from tables and figures using WebPlotDigitizer software version 4.3 (Rohatgi, 2020). A full list of data extracted is found in Table S2.

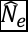

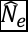

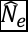

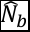

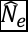

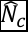

Table S1 – Single-sample genetic *N_e_* estimation methods

| Estimation Method | Description | Software | Reference |
| --- | --- | --- | --- |
| Approximate Bayesian Computation | Uses summary statistics from a population with Bayesian computation to provide posterior probability distributions for N_e_. | ONeSAMP | (Tallmon, Koyuk, Luikart, & Beaumont, 2008) |
| Heterozygote Excess | Estimates the effective number of breeders based on the assumption that when N_b_ is small, there will be an excess of heterozygotes (Pudovkin et al., 1996; Zhadnova & Pudovkin, 2008). | NeEstimator v2 | (Do et al., 2014) |
| Linkage Disequilibrium (LD) | Uses non-random associations of alleles at different loci (i.e., deviations from the expected genotype frequency based on random distribution) to estimate random drift and therefore N_e_ or N_b_ (Hill, 1980). A bias correction by Waples (2006) eliminates downward bias when sample size is less than the true N_e_. | LDNe | (Waples & Do, 2008) |
|  |  | NeEstimator v2 | (Do et al., 2014) |
| Molecular Coancestry | Estimates the effective number of breeders from molecular coancestry, a measure of shared alleles between individuals (Nomura, 2008). | NeEstimator v2 | (Do et al., 2014) |
| Sibship Frequency | Estimates demographic parameters from genotypes of a cohort, and then uses these with predictive equations based on the probabilities of sampling half-sibs or full-sibs from the population (Wang, 2009). | COLONY | (Jones & Wang, 2010) |

Table S2 – Data extraction protocol used by reviewers

| **Group** | **Column** | **Explanation** |
| --- | --- | --- |
| **Population information** | Common Name | Common name of species, used by authors in article, all lower case |
|  | Genus | genus (capitalized) |
|  | Species | species (uncapitalized) |
|  | Taxonomic group | freshwater fish, marine fish, diadromous fish, reptile, amphibian, mammal, bird, invertebrate, plant. If something is not listed here, or you are unsure, you can enter a comment in the column next to this. For fish that can be either resident or diadromous, use best judgement according to the authors’ descriptions of the population |
|  | Population | name given to population; for species in bodies of water, could use the name of the lake/river, or another name used by authors in article |
|  | Population ID | numerical identification for each population (since there can be multiple estimates for a single population). Alpha-numeric system using the article number. i.e. if article # 100 has two populations, they will be 100A and 100B. If an article only has one estimate, still include A at the end. |
|  | Method of defining population | based off of the authors in the article and how they defined the population. E.g. using Fst values, STRUCTURE (determining # of groups), BAYESASS (measuring migration rates), IBA (individual-based-assignment; using genetic data from populations to assign individuals), etc. If there is any additional information, include it in the comment column. E.g. what their threshold Fst value was, or level of migration, etc. |
|  | Location (coordinates) | coordinates in lat/long, decimal degrees, to 6 decimals if available. If article has info on UTM, use this website to convert: https://www.engineeringtoolbox.com/utm-latitude-longitude-d_1370.html . If article has info on lat/long in degrees/minutes/seconds, use this website to convert: https://www.latlong.net/degrees-minutes-seconds-to-decimal-degrees . If article includes a map or description of location but does not provide coordinates, use google maps to generate coordinates as accurately as possible (e.g. if given the name of a lake, take the coordinates in middle of lake). You can also use google maps to find the "average" coordinates if you are pooling sampling locations that make up a single population (helpful tool is "measure distance" and can take the midpoint) |
|  | Region where population is located | can be a city/province/ etc. or multiple of these things. |
|  | Country | free-form text. Please capitalize country name, and use as accurate spelling as possible. If samples were taken from multiple countries, you can enter one here, and add the others in the "region" column. |
|  | Continent/Ocean | drop-down list of continents and oceans. For anything not included in the drop-down list, you can enter it in the "region" column. |
|  | Was the pop stocked or supplemented? | e.g. historical supplementation, stocking into a new location, re-introductions (that are now reproducing independently) |
|  | Is the pop non-native? | non-native or invasive species |
|  | When was the pop stocked/ introduced? | the year when pop was introduced or last stocked |
|  | notes | notes on population information |
| **Protection status** | At-risk | YES/NO (based on information in the article) |
|  | Protection status | threatened, vulnerable, endangered, etc. (enter "other" into comments column). Based on what is written in article by authors |
|  | Protection status organization | e.g. IUCN, COSEWIC, USFW, etc. (population-specific info is better if mentioned, instead of IUCN) |
| **Method** | Method (general) | LD (linkage disequilibium), SF (sibship frequency), HE (heterozygote excess), MC (molecular coancestry), Bayesian methods. If article used a different single-sample genetic estimator not listed, put "Other" and enter it into the notes column |
|  | Method (specific) | LDNe, NeEstimator (v1 or V2), COLONY, ONeSAMP. If article used a different software not listed, put "Other" and enter it into the notes column |
|  | Marker type | microsatellite, allozyme, SNP, etc. |
|  | GW correction | genome-wide bias correction for LD method; YES/NO. Based on study by Waples, Larson, and Waples (2016, Heredity) |
|  | Allele freq cutoff | for LD method; common values are 0.01, 0.05, 0.1. If the article reports multiple allele cutoff values, follow this rule: For sample sizes >25 use 0.02, and <25 use 0.05. Please report whether the study included multiple allele cutoffs. |
|  | Mating system | for SF method. If the mating system is not one of the given options, please choose "other" and enter it in the comments column. If the study includes data on multiple mating systems, choose the best option based on the authors' discussion, and make a comment that there were multiple values reported |
|  | other comments for SF method | here, please include any other details about the SF method that the authors report. E.g. whether it was based on random mating or inbreeding; the probability of an offspring in a dataset having a parent in the dataset; etc. If multiple values are given for any of these, choose one either based on author discussion, or choose the median value (e.g. if chance of sampling a parent is reported at 0.3, 0.5 and 0.7, use 0.5), and make a comment that other values are reported in the article |
|  | Priors | For the Bayesian method in ONeSAMP. Upper and lower boundaries of the Ne made prior to estimation. (e.g. 5-150). If multiple values are given, choose the estimate with the widest range of priors. (i.e. choose 2-50 rather than 10-20, etc.) |
|  | Type of sequencing | RAD-seq, GBS, capillary electrophoresis, Sanger. Can enter a method not listed here by choosing "other" and then entering in comment column. |
|  | # of loci/SNP | number of loci or SNP used in estimate. **ensure you adjust this number if the study excludes monomorphic loci or loci with high null freq.** |
|  | He and Ho | average He (expected heterozygosity) or Ho (observed heterozygosity) across loci for that population. For microsatellites. |
|  | Ar | allelic richness; average # of alleles per locus, weighted by sample size. If the authors refer to "Ar" with no mention of method, assume they are correct. If they refer to Ar as the non-weighted version, then please categorize as MNA instead. |
|  | MNA | mean number of alleles per locus. NOT weighted. If the authors refer to MNA but mention weighting, categorize as Ar instead. If they refer to MNA with no mention of method, assume they are correct. |
|  | Inbreeding coefficient (Fis) | usually calculated from heterozygosity measures. |
|  | Nucleotide diversity | for SNPs |
|  | Sample size | the number of individuals sampled from the population |
| **Ne values** | Ne | point estimate of Ne. If the value is infinite, enter "infinity" (please record these for now; though they may be excluded later) |
|  | Nb | point estimate of Nb |
|  | Year estimate was taken | year the samples were taken from the population. If no year is given, leave blank. If multiple years, use most recent year but make note of other years. |
|  | LCI | lower confidence interval for estimate (can be "infinity" as well) |
|  | UCI | upper confidence interval for estimate (can be "infinity" as well) |
|  | CI method | method used to calculate Cis. E.g. jackknife vs parametric methods in LDNe program. If the method is not provided, enter as text in comments column. |
| **Ne verification** | Did they sample across cohorts? | yes/no/unsure. in order to be an accurate estimate of Ne, there need to be sampling across different cohorts (i.e. birth years). If only one cohort is sampled, then this is Nb, not Ne. |
|  | Did they report Ne for sampling sites? | If the authors define a population as a group of sampling sites, but only report Ne for the sites (rather than for the population as a whole), please mark this column as "YES", and report each sampling site on a unique row, with the SAME POPULATION ID. If Ne is reported for both the sampling sites, and overall population, use the population-level Ne, and mark this column as NO, because you are reporting on the population level. |
|  | Did they pool samples from multiple years? | yes/no/unsure. Put other years in the notes column |
|  | Notes | any other notes on the validity of the Ne estimate (e.g. all samples came from a single breeding site and may not be representative of the population) |
| **Nc values** | Nc estimate and Cis | same as Ne, but with Nc |
|  | Nc method | mark-recapture (in comment, please put the type of mark-recapture. i.e. petersen/lincoln-petersen, schnabel, etc.), complete count, incomplete count (e.g. quadrat study with extrapolation). If the method is not in the list, please put "other" and then write out method in comment column. |

Table S3 – Replication within the database

| **Type of Replication** | **Description** | **Which records were retained?** |
| --- | --- | --- |
| Marker Replication | For a single population, the authors used both microsatellites and SNPs to estimate *N_e_* or *N_b_*. | SNP estimates, as they had a higher power (which we calculated by multiplying the number of markers, sample size, and allelic richness) |
| Software Replication | Authors estimated N_e_ or N_b_ using both NeEstimator V2 and LDNe | Estimates generated in NeEstimator V2, as it is the more recent software, and includes an updated bias correction to address missing data |
| Spatial Replication | Multiple estimates from a single population (i.e. they all shared the same population ID), for example when N_e_ was reported for several sampling sites that were shown through Fst or STRUCTURE to cluster into one population | The estimate with the largest sample size (S), as larger sample sizes generally produce more reliable estimates. In the case where two estimates with the same population ID had the same sample size, we chose either the estimate with a non-infinite upper confidence interval, or we chose randomly (using a random number generator) if all upper CIs were finite. |
| Temporal Replication | Multiple estimates were reported over time for the same population | The most recent estimate, as these were most relevant for a contemporary review of N_e_ |

Table S4 – Backward selection of Generalized Linear Mixed Models using AICc. Bold indicates the most parsimonious model, and * indicates interactions. The random effects are indicated in parentheses, as are all random intercept effects.

| Analysis | Equation* | df | AICc | ΔAICc |
| --- | --- | --- | --- | --- |
| $\hat{N}_{e}$ 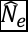 | N_e_ ~ taxonomic group + marker + loci number + (1 \| StudyID / PopulationID) | 15 | 3121281 | 0.000 |
|  | **N_e_ ~ taxonomic group + marker + (1 \| StudyID / PopulationID)** | 14 | 3121280 | –1.721 |
| $\hat{N}_{b}$ 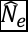 | N_b_ ~ taxonomic group + marker + loci number + (1 \| StudyID / PopulationID) | 14 | 970305.9 | 0.000 |
|  | **N_b_ ~ taxonomic group + marker + (1 \| StudyID / PopulationID)** | 13 | 970304.7 | –1.208 |
| Ratio | Ratio ~ taxonomic group + marker + loci number + ratio type + (1 \| StudyID / PopulationID) | 15 | –102471.7 | 0.000 |
|  | Ratio ~ taxonomic group + loci number + ratio type + (1 \| StudyID / PopulationID) | 14 | –102473.8 | –2.106 |
|  | **Ratio ~ taxonomic group + ratio type + (1 \| StudyID / PopulationID)** | 13 | –102472.8 | –1.049 |
| Fifty | Fifty ~ taxonomic group + marker + loci number + (1 \| StudyID) | 12 | 3602.6 | 0.000 |
|  | Fifty ~ taxonomic group + marker + (1 \| StudyID) | 11 | 3602.1 | –0.452 |
|  | **Fifty ~ taxonomic group + (1 \| StudyID)** | 10 | 3602.8 | 0.215 |
| Fivehundred | Fivehundred ~ taxonomic group + marker + loci number + (1 \| StudyID) | 12 | 2205.3 | 0.000 |
|  | **Fivehundred ~ taxonomic group + marker + (1 \| StudyID)** | 11 | 2203.3 | –2.014 |
| IUCN | N_e_ ~ taxonomic group * IUCN + marker + loci number + (1 \| StudyID / PopulationID) | 24 | 2291542 | 0.000 |
|  | **N_e_ ~ taxonomic group + IUCN + marker + loci number + (1 \| StudyID / PopulationID)** | 16 | 2291540 | -1.472 |
| HFI | N_e_ ~ taxonomic group * HFI + marker + loci number + (1 \| StudyID / PopulationID) | 24 | 2004509 | 0.000 |
|  | **N_e_ ~ taxonomic group * HFI + marker + (1 \| StudyID / PopulationID)** | 23 | 2004507 | –2.024 |

Table S5 – Sample sizes by taxonomic group for all analyses. Ratio = $\hat{N}_{e}$*/*$\hat{N}_{c}$ or $\hat{N}_{b}$*/*$\hat{N}_{c}$

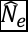

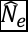

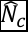

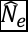

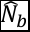

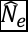

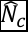

| **Taxonomic Group** | $\hat{N}_{e}$ **Analysis** 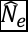 | $\hat{N}_{b}$ **Analysis** 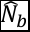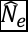 | **50/500 Analyses** | **IUCN Analysis** | **Human Footprint Analysis** | **Ratio Analysis** |
| --- | --- | --- | --- | --- | --- | --- |
| Amphibians | 369 | 68 | 350 | 368 | 339 | 25 |
| Diadromous Fishes | 686 | 213 | 571 | 536 | 506 | 107 |
| Birds | 204 | 9 | 145 | 204 | 165 | 41 |
| Freshwater Fishes | 1113 | 360 | 860 | 656 | 1045 | 140 |
| Invertebrates | 321 | 60 | 253 | 129 | 216 | 8 |
| Mammals | 552 | 69 | 440 | 544 | 477 | 96 |
| Marine Fishes | 248 | 33 | 178 | 197 | 22 | 9 |
| Plants | 439 | 5 | 350 | 338 | 387 | 80 |
| Reptiles | 201 | 10 | 191 | 200 | 132 | 33 |
| **Total** | **4145** | **827** | **4145** | **3172** | **3299** | **537** |

Table S6 – Median *N_e_* and *N_b_* estimates across taxonomic groups

| **Taxonomic Group** | **Median** $\hat{N}_{e}$ 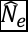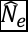 | **Median** $\hat{N}_{b}$ 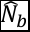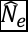 | **Median Ratio between** $\hat{N}_{e}$ **and** $\hat{N}_{c}$ 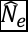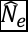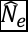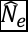 |
| --- | --- | --- | --- |
| Amphibians | 43.3 | 52.15 | 0.13 |
| Diadromous Fishes | 211.5 | 113.8 | 0.40 |
| Birds | 77.8 | 71.0 | 0.48 |
| Freshwater Fishes | 80.8 | 75.25 | 0.12 |
| Invertebrates | 135.6 | 173.2 | 0.09 |
| Mammals | 47.2 | 93.0 | 0.17 |
| Marine Fishes | 634.6 | 798.0 | 0.45 |
| Plants | 35.1 | 402.6 | 0.16 |
| Reptiles | 72.3 | 40.65 | 0.14 |
| **Overall** | **83.9** | **93.0** | **0.24** |

Table S7 – Sample sizes and percentages by taxonomic group for the IUCN Red List designations. “Threatened” = Vulnerable, Endangered, Critically Endangered. “Nonthreatened” = Least Concern, Near Threatened. “N/A” = Data Deficient, Not Evaluated.

| Taxonomic Group | Threatened | | Nonthreatened | | N/A | |
| --- | --- | --- | --- | --- | --- | --- |
|  | N | Percentage | N | Percentage | N | Percentage |
| Amphibians | 67 | 15.3 | 369 | 84.4 | 1 | 0.2 |
| Diadromous Fishes | 98 | 10.9 | 619 | 68.8 | 182 | 20.2 |
| Birds | 47 | 22.1 | 166 | 77.9 | 0 | 0 |
| Freshwater Fishes | 216 | 14.7 | 646 | 43.9 | 611 | 41.5 |
| Invertebrates | 53 | 13.9 | 80 | 21.0 | 248 | 65.1 |
| Mammals | 169 | 27.2 | 441 | 71.0 | 11 | 1.8 |
| Marine Fishes | 95 | 33.8 | 126 | 44.8 | 60 | 21.3 |
| Plants | 99 | 22.3 | 244 | 54.9 | 101 | 22.7 |
| Reptiles | 103 | 48.8 | 107 | 50.7 | 1 | 0.5 |
| Total | 947 | 19.1 | 2798 | 56.4 | 1215 | 24.5 |

Table S8 – Parameter estimates from Generalized Linear Mixed Model comparing $\hat{N}_{e}$ to Human Footprint Index across taxonomic groups. Estimates are shown on the log-scale

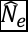

| **Taxonomic Group** | **Slope** | | **Intercept** | |
| --- | --- | --- | --- | --- |
|  | Estimate | SE | Estimate | SE |
| Amphibians | -0.0238 | 0.0019 | 4.91 | 0.37 |
| Diadromous Fishes | 0.0059 | 0.0008 | 5.46 | 0.34 |
| Birds | -0.0200 | 0.0016 | 5.58 | 0.35 |
| Freshwater Fishes | -0.0127 | 0.0005 | 5.26 | 0.31 |
| Invertebrates | -0.0178 | 0.0013 | 5.72 | 0.32 |
| Mammals | -0.0185 | 0.0016 | 4.64 | 0.31 |
| Marine Fishes | -0.0047 | 0.0165 | 6.65 | 0.82 |
| Plants | 0.0199 | 0.0014 | 2.95 | 0.32 |
| Reptiles | -0.0092 | 0.0059 | 4.82 | 0.43 |

Table S9 – Number of populations per taxonomic group with an upper confidence interval (CI) of infinity

| **Taxonomic Group** | **Populations with  Infinitive upper CI (N)** | **Percentage (%)** | **Total N** |
| --- | --- | --- | --- |
| Amphibians | 136 | 31.12 | 437 |
| Diadromous Fishes | 198 | 22.02 | 899 |
| Birds | 55 | 25.82 | 213 |
| Freshwater Fishes | 459 | 31.16 | 1473 |
| Invertebrates | 191 | 50.13 | 381 |
| Mammals | 83 | 13.36 | 621 |
| Marine Fishes | 109 | 38.79 | 281 |
| Plants | 80 | 18.01 | 444 |
| Reptiles | 47 | 22.27 | 211 |

Table S10 – Average and Variance of sample sizes per taxonomic group, using the full dataset.

| **Taxonomic Group** | **Average Sample Size** | **Median Sample Size** | **Standard Deviation of Sample Size** | **Minimum Sample Size** | **Maximum Sample Size** |
| --- | --- | --- | --- | --- | --- |
| Amphibians | 33.69 | 27 | 54.46 | 4 | 982 |
| Diadromous Fishes | 86.03 | 58 | 145.17 | 6 | 2577 |
| Birds | 62.42 | 35 | 68.45 | 5 | 422 |
| Freshwater Fishes | 71.37 | 41 | 171.21 | 5 | 3805 |
| Invertebrates | 58.97 | 31 | 227.7 | 6 | 4041 |
| Mammals | 61.36 | 34 | 85.59 | 6 | 1175 |
| Marine Fishes | 130.51 | 52 | 433.16 | 7 | 5413 |
| Plants | 40.16 | 30 | 46.36 | 4 | 418 |
| Reptiles | 46.58 | 27 | 66.32 | 5 | 666 |
| Total | 67.62 | 39 | 171.26 | 4 | 5413 |

**Identification of studies via databases and registers**

Records removed *before screening*:

Duplicate records removed
(n = 660)

Records identified from*:

WebOfScience (n = 3875)

LDNe search (n = 798)

NeEstimator search (n = 500)

**Identification**

Studies excluded

(n = 2016)

Studies screened at Title and Abstract (n = 4513)

Studies not retrieved

(n = 32)

Studies sought for retrieval

(n = 2497)

**Screening**

Studies excluded:

No *N_e_* estimate (n = 1105)

Temporal method (n = 251)

Supplemented pop (n = 62)

Human/virus/pathogen (n = 68)

Mitochondrial DNA (n = 9)

Review article (n = 13)

Duplicate articles (n = 29)

Studies screened at Full Text

(n = 2465)

Studies included in initial database

(n = 928)

Estimates of *N_e_* from included studies

(n = 8971)

**Included**

Fig. S1: Flow diagram of the review process, modified from O’Dea et al., 2021

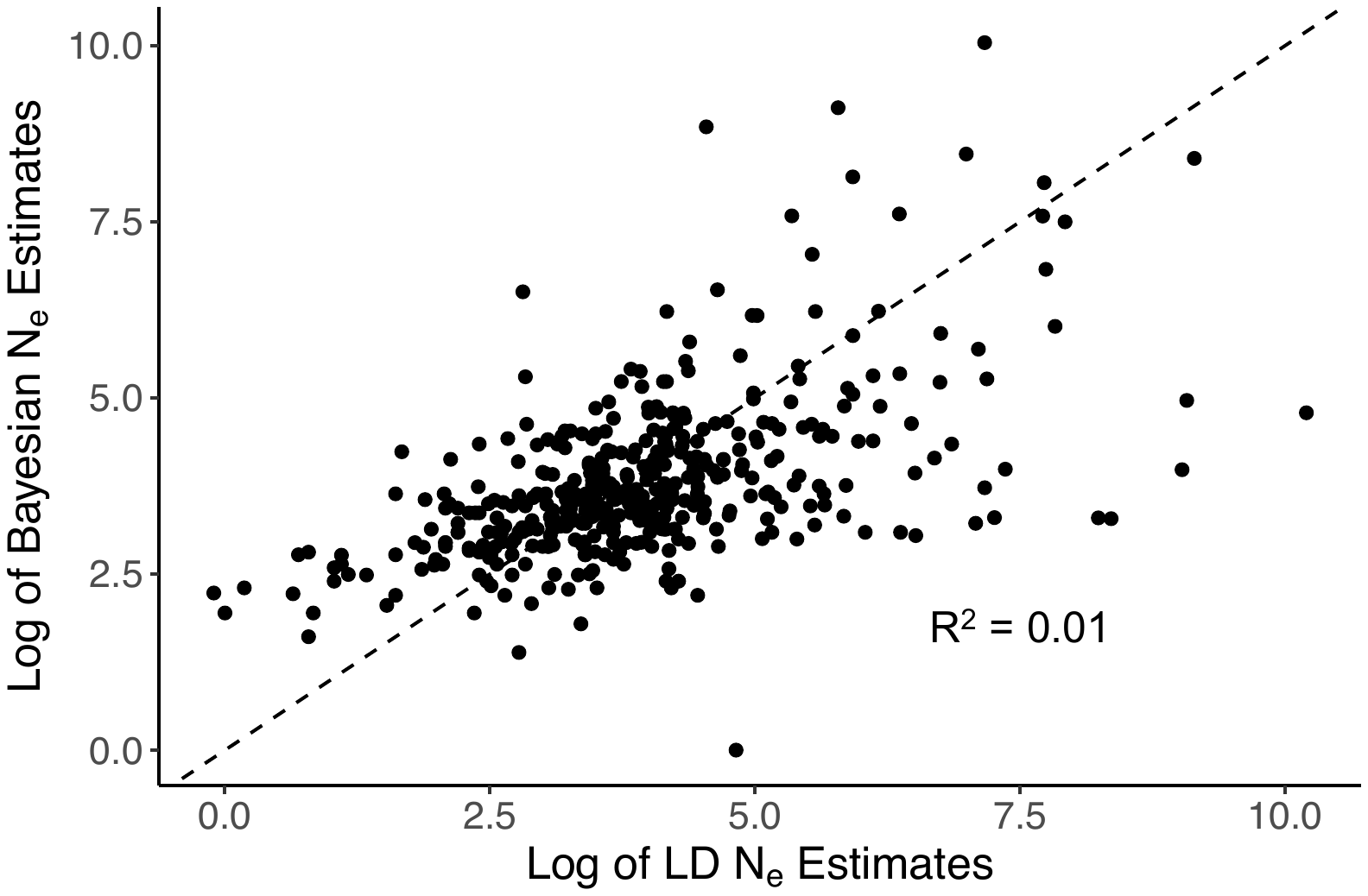

Fig. S2: Regression between *N_e_* estimates produced from the Linkage Disequilibrium (LD) method, and the Bayesian Computation method. The dataset is made up of estimates from studies that used both the LD and Bayesian estimation methods on the same populations, from the same time periods. Sample size for the analysis was N = 416.

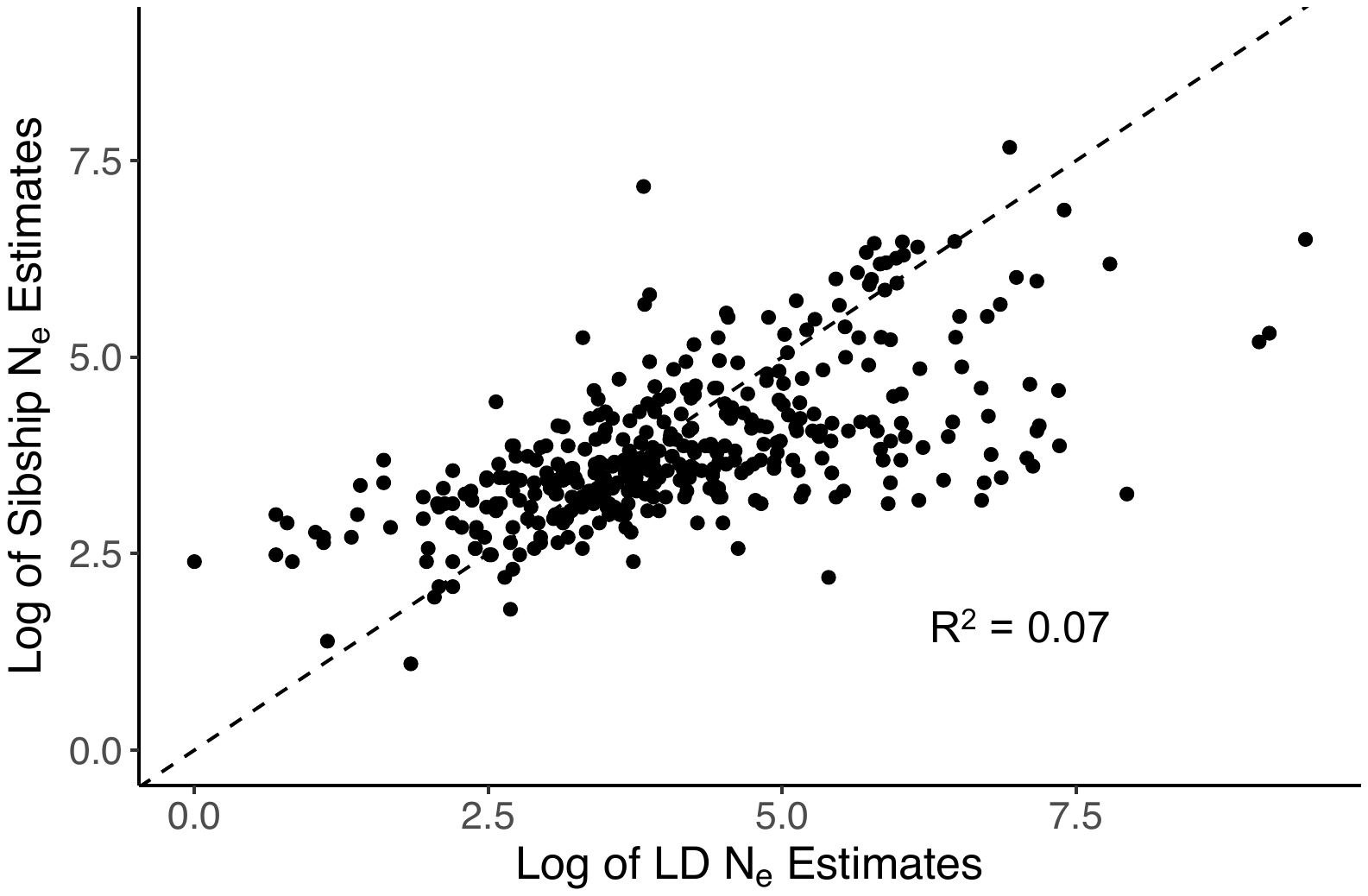

Fig. S3: Regression between *N_e_* estimates produced from the Linkage Disequilibrium (LD) method, and the Sibship Frequency method. The dataset is made up of estimates from studies that used both the LD and Sibship estimation methods on the same populations, from the same time periods. Sample size for the analysis was N = 386.

**Filtering of Data**

Records removed *based on method*:

Non-Linkage Disequilibrium

(n = 2347)

Estimates generated from:

Linkage Disequilibrium

(n = 6624)

Sibship Frequency (n = 928)

Bayesian (n = 850)

Other (n = 569)

**Estimation Method**

Records removed:

Infinite or negative estimates

(n = 854)

No bias correction (n = 247)

Duplicate populations (n = 126)

No sample size (n = 57)

No point estimate (n = 50)

Only one sex sampled (n = 15)

Did not meet screening

requirements (n = 218)

Estimates screened from Linkage Disequilibrium Method (n = 6624)

**Filtering**

Records removed *based on replication*:

Marker replication (n = 74)

Software replication (n = 11)

Estimates after filtering
(n = 5057)

**Replication**

Records removed *from dataset without replication*:

Temporal replication (n = 789)*

Spatial replication (n = 606)*

Final dataset (n = 4972)

Final dataset without replication (n = 3658)

**Included**

Fig. S4: Flow diagram of the filtering process, modified from O’Dea et al., 2021. *Some records had both spatial and temporal replication

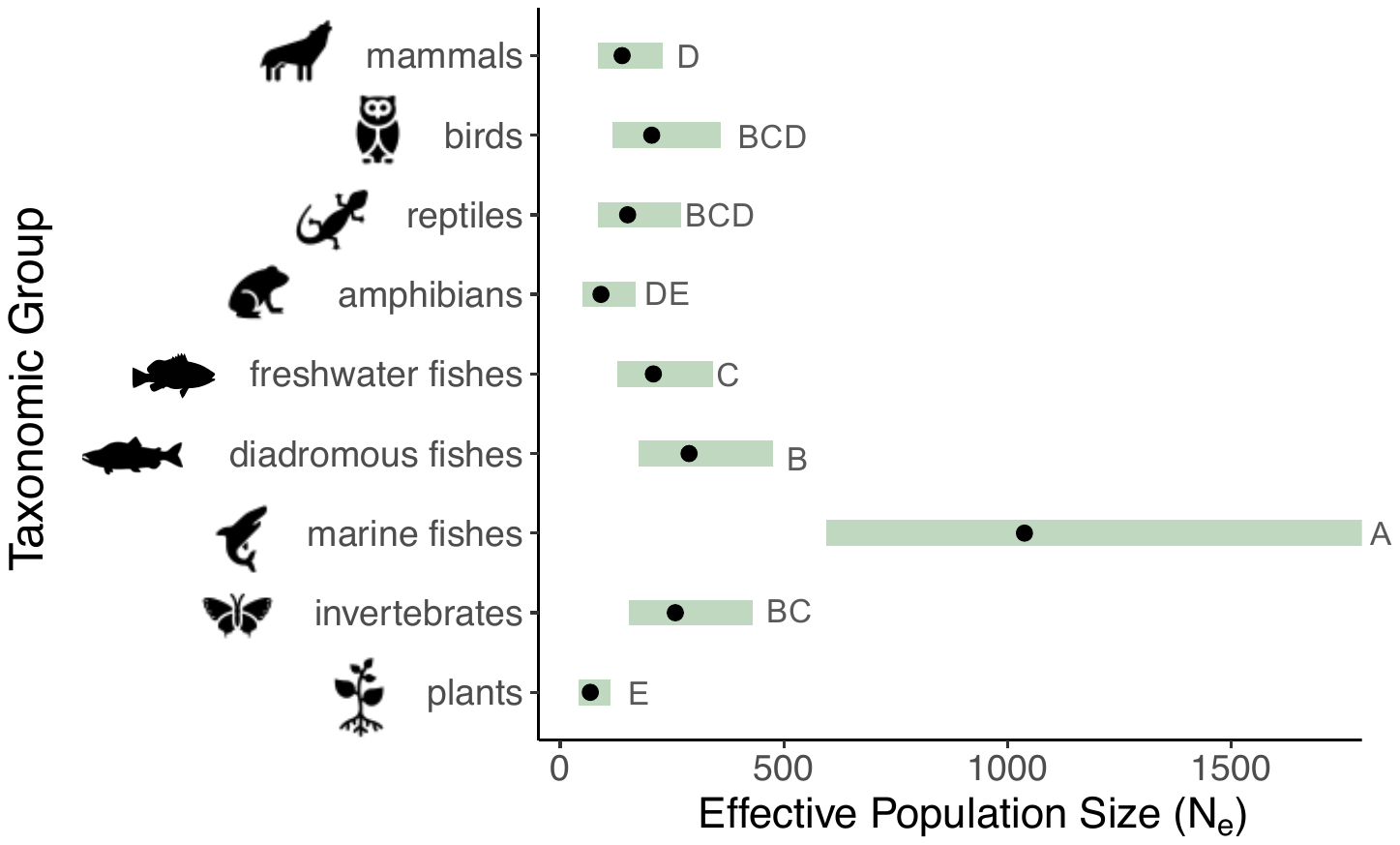

Fig. S5: Mean effective population size ($\hat{N}_{e}$) across taxonomic groups, when accounting for populations with an upper CI of infinity. Shaded green bars represent 95% confidence intervals. Groups with no shared letters are statistically different from one another

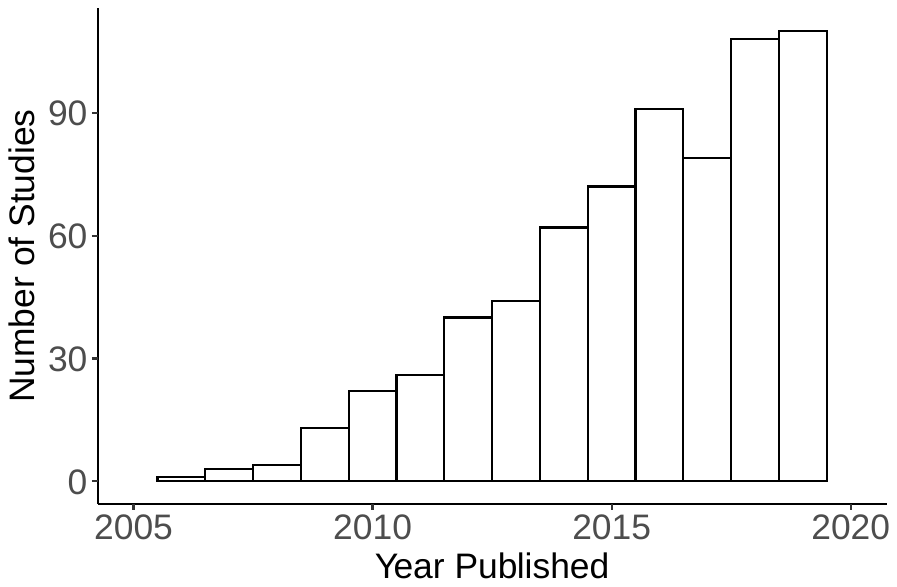

Fig. S6: Number of studies with *N_e_* or *N_b_* estimates published per year. Studies from 2020 were not included as the literature search was conducted partway through the year and is not representative of all articles published

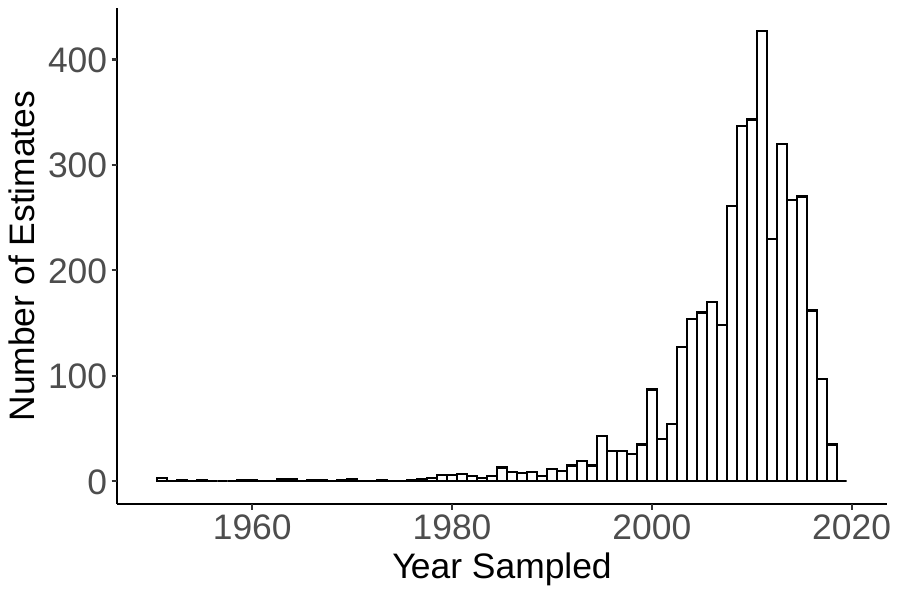

Fig. S7: Number of estimates of *N_e_* or *N_b_* sampled per year in the final dataset. There were 957 estimates with no sampling year reported.

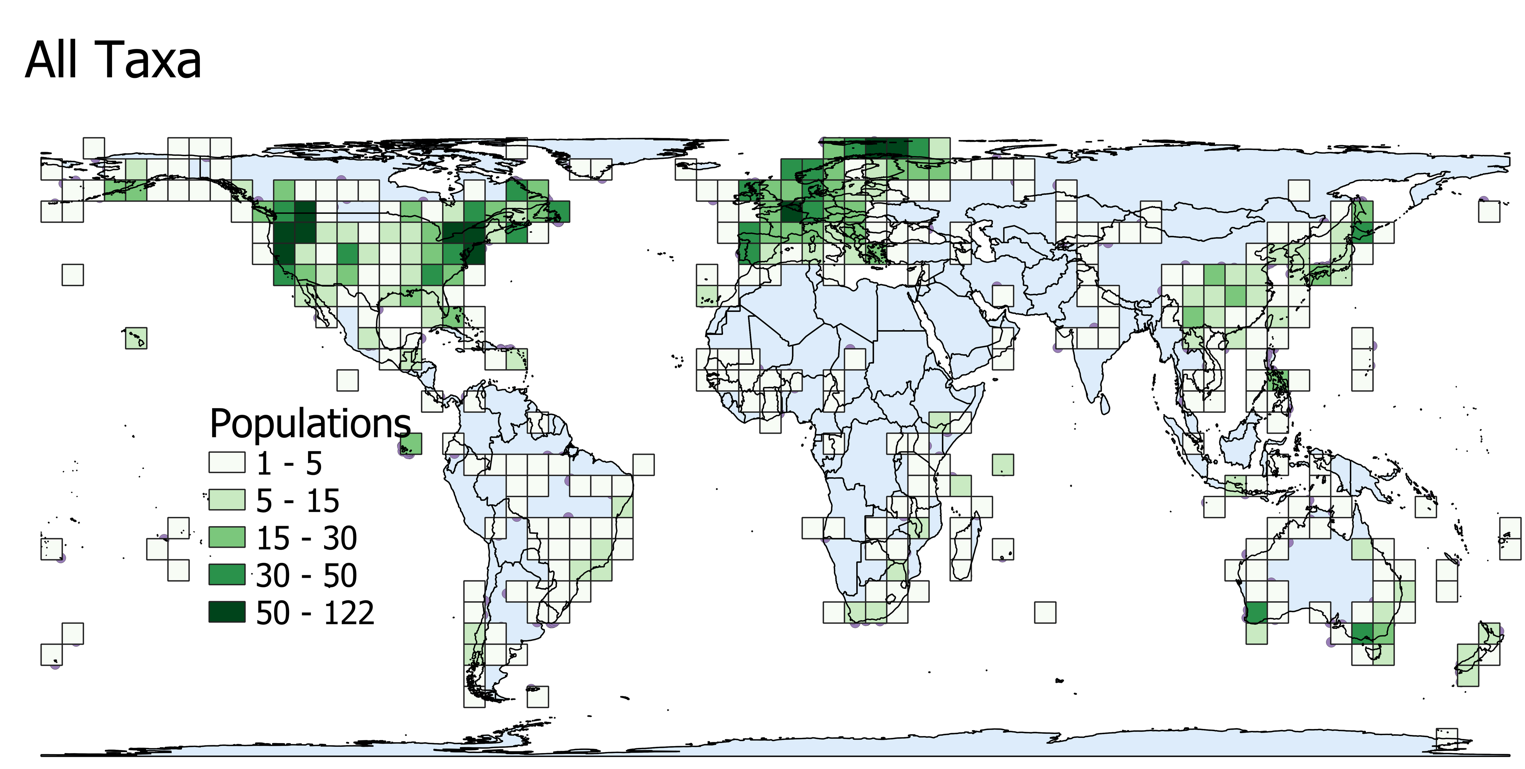

Fig. S8: Global map showing the number of populations sampled in each 250,000 km^2^ grid cell. Data is projected with the world Behrmann projection

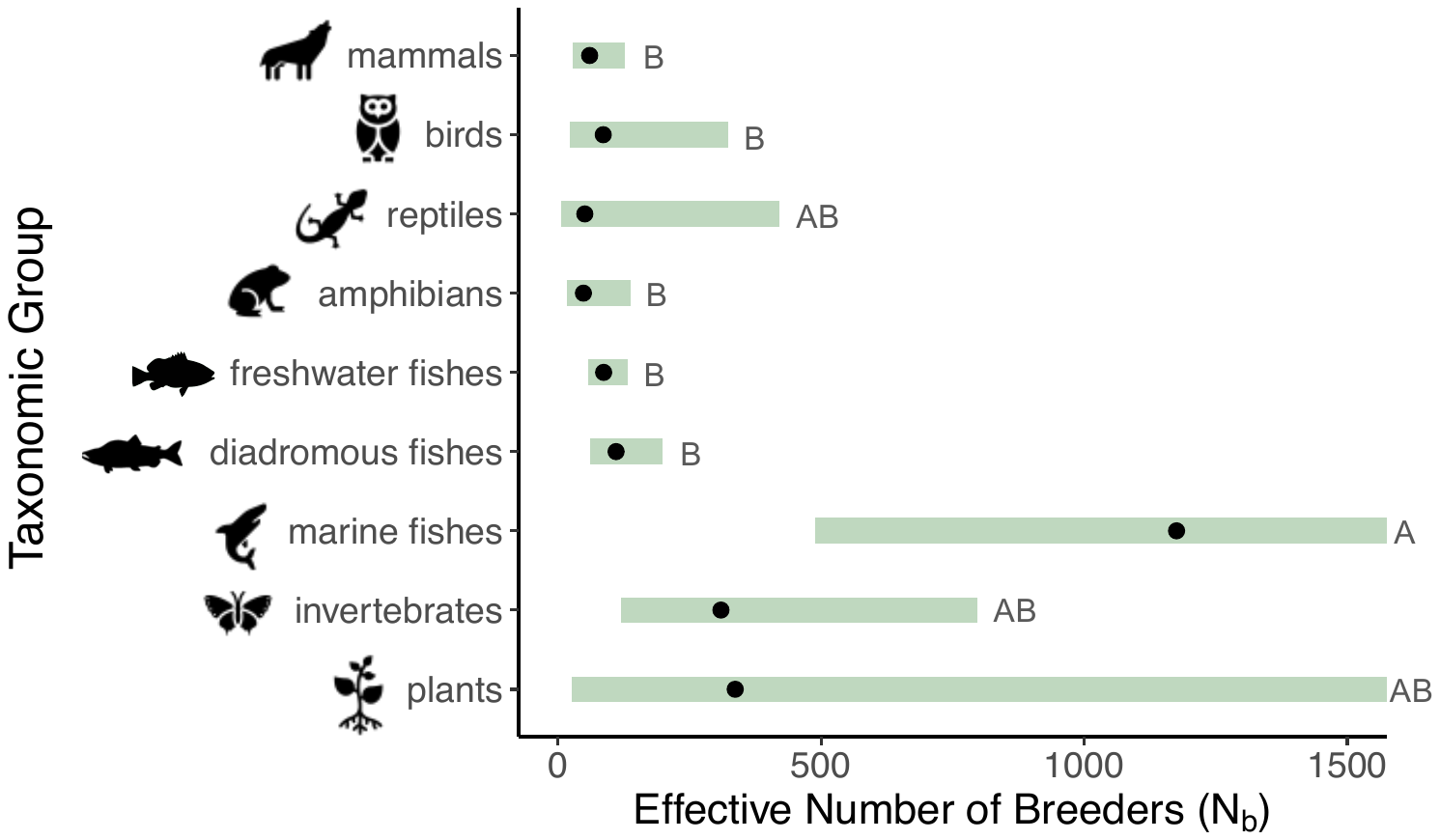

Fig. S9: Mean effective number of breeders ($\hat{N}_{b}$) across taxonomic groups, accounting for differences in marker type. Shaded green bars represent 95% confidence intervals. *The upper CI for plants and marine fish are not contained within the plot. Groups with no shared letters are statistically different from one another

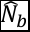

Fig. S10: Global maps showing the taxa-specific median $\hat{N}_{e}$ value in each 250,000 km^2^ grid cell. Data is projected with the world Behrmann projection

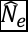
